# Domains Required for FRQ-WCC Interaction within the Core Circadian Clock of *Neurospora*

**DOI:** 10.1101/2023.02.25.530043

**Authors:** Bin Wang, Jay C. Dunlap

## Abstract

In the negative feedback loop composing the *Neurospora* circadian clock, the core element, FREQUENCY (FRQ) binds with FRH (FRQ-interacting RNA helicase) and Casein Kinase 1 (CK1) to form the FRQ-FRH complex (FFC) which represses its own expression by interacting with and promoting phosphorylation of its transcriptional activators White Collar-1 (WC-1) and WC-2 (together forming the White Collar Complex, WCC). Physical interaction between FFC and WCC is a prerequisite for the repressive phosphorylations, and although the motif on WCC needed for this interaction is known, the reciprocal recognition motif(s) on FRQ remains poorly defined. To address this, FFC-WCC was assessed in a series of *frq* segmental-deletion mutants, confirming that multiple dispersed regions on FRQ are necessary for its interaction with WCC. Biochemical analysis shows that interaction between FFC and WCC but not within FFC or WCC can be disrupted by high salt, suggesting that electrostatic forces drive the association of the two complexes. As a basic sequence on WC-1 was previously identified as a key motif for WCC-FFC assembly, our mutagenetic analysis targeted negatively charged residues of FRQ leading to identification of three Asp/Glu clusters in FRQ that are indispensable for FFC-WCC formation. Surprisingly, in several *frq* Asp/Glu-to-Ala mutants that vastly diminish FFC-WCC interaction, the core clock still oscillates robustly with an essentially WT period, indicating that the binding strength between the positive and negative elements in the feedback loop is required for the clock but is not a determinant of the period length.

## Introduction

Circadian rhythms control a wide range of cellular and behavioral processes in most eukaryotes and certain prokaryotes (1, 2), facilitating adaptation of life to constant environmental changes; disruption of rhythms has been implicated in various diseases in humans (3). At the molecular level, circadian clocks rely mainly on interlocked positive and negative arms, and in circadian cycles the latter gradually repress their own expression through inactivating the former on a timescale of hours. In the clock of *Neurospora crassa*, a circadian model organism used for decades, the White Collar Complex (WCC) derived from all cellular WC-1 and a fraction of WC-2 drives transcription of the central pacemaker gene, *frequency (frq)*, through binding to either of two DNA elements in the *frq* promoter under contrasting illumination conditions either the *Clock box (C-box)* in the dark or the *Proximal Light-Response Element (pLRE)* upon light exposure or in constant light (4–6). FREQUENCY (FRQ), encoded by the frq gene, associates with FRH (FRQ-interacting RNA helicase) (7, 8) and CKI (Casein Kinase I) (9) to create the FFC (FRQ-FRH complex), repressing WCC transcriptional activity by promoting its phosphorylation at a group of residues (9–11) and thereby terminating its own expression to conclude a circadian cycle.

An orthologous feedback loop is found in mammalian cells, with CLOCK and BMAL1 forming a heterodimer that plays the role of WCC, driving expression of PERs and CRYs which form a negative element complex with CK1, analogous to the FFC, that inactivates CLOCK/BMAL1 by phosphorylation at key residues (12, 13). As is the case in *Neurospora*, while much is becoming known about the means through which repression is achieved (13–17), relatively little is known about the structural determinants on PER/CRY or CLOCK/BMAL that determine their interactions.

In *Neurospora*, WCC acts as a responsive hub coordinating light signaling and the circadian clock that is obligate in constant darkness (18). In consonance with its dual role as a light sensor and a transcription factor for frq in the clock, WC-1 bears a light-, oxygen-, and voltage-sensing (LOV) domain, a transactivation domain, two motifs required for DNA binding—the zinc-finger (ZnF) and its nearby DBD (defective in DNA binding) motif, and two Per-Arnt-Sim (PAS) domains for WC-2 interaction (19); WC-2, an accessory of WC-1, contains a PAS and a ZnF DNA-binding domain (19). The DBD motif (KKKRKRRK) on WC-1 has been discovered to be indispensable for WCC to bind the *C box* and drive *frq* transcription in the dark as well as for interacting with FRQ/FRH (20).

FRQ functions mainly as an organizing platform for recruiting and scaffolding the circadian negative elements FRH and CKI in forming the FRQ-FRH-CKI complex which controls the pace of the clock (7, 21). Self-interaction among FRQ molecules occurs via their coiled-coil (CC) domains, and disruption of the CC results in arrhythmicity (22). The nuclear localization signal (NLS) of FRQ is necessary for its circadian function in WCC repression (23). FRQ interacts with CKI by two domains: FRQ-CK1a interaction domain 1 (FCD1) (24) and FCD2 (9) and with FRH via the FFD (FRQ–FRH interaction Domain) (7, 25). FRQ also contains two PEST-like elements: PEST-1 and PEST-2, both of which undergo CK-1a- and CK-1b-mediated progressive phosphorylations, and elimination of PEST-1 results in arrhythmic conidiation, reflecting an impaired clock (26). Intramolecular interplay between the N- and C-termini of FRQ has been noticed as key for the clock operation (24).

Multisite phosphorylation has been investigated extensively as a major mechanism finely tuning circadian activities of both FRQ (27–30) and its transcriptional factor WCC (9–11, 31). As the initial step in feedback-loop execution, negative elements need to interact physically with the positive factors in order to bring about timely but gradual repression on the latter. While prior work on the *Neurospora* clock has focused on phosphorylation and transcription-centered mechanisms controlling the clock, it remains largely unknown how molecular contacts between the two limbs are accomplished, though this step is plainly required for subsequent negative feedback inhibition. We previously uncovered a motif on WC-1 composed of eight consecutive highly basic residues required for interaction with FRQ (20), raising a corresponding query regarding which regions and residues on FRQ are reciprocally involved in the establishment of FFC-WCC. To tackle this puzzle, here we have characterized key regions on FRQ essential for the WCC incorporation and the core oscillator. As biochemical data hinted that FFC and WCC may contact via electrostatic charges, we focused on negatively charged residues falling in the important regions of FRQ that determine WCC interaction. To this end, mutagenetic analyses unearthed three clusters of residues on FRQ that contribute to WCC association. Interestingly however, the core clock still runs normally in certain *frq* mutants with an extremely low level of FFC-WCC, revealing that while interaction between the positive and negative arms is necessary for the feedback loop, it is not a rate-limiting step for period length.

## Results

### A *frq* deletion series identifies regions required for the *Neurospora* clock

Three distinct regions on FRQ have been implicated in WCC interaction: amino acids 107-310, 435-558, and 631-905 (25). The loss of FRQ-WCC interaction in *frq* mutants missing any of these pieces (25) might be attributed to the removal of the WCC-interacting domain(s) or pleiotropic effects caused by polypeptide shortening. To address this, we engineered a set of *frq* mutants each missing ∼50 amino acids situated in these regions. In these mutants, *frq* transcription as reported by a C-box-driven luciferase gene was measured in real-time (Figure 1). *frq^Δ564-603^* shows a WT period length; *frq^Δ691-732^* displays a shortened period of 15.5 hrs; *frq^Δ107-148^*, *frq^Δ435-481^*, *frq^Δ604-646^*, *frq^Δ874-884^*, and *frq^Δ857-905^* have prolonged rhythms to variable extents. The core clock became totally arrhythmic in *frq^Δ149-193^, frq^Δ194-199^, frq^Δ200-249^, frq^Δ250-310^, frq^Δ482-510^, frq^Δ511-558^, frq^Δ647-690^, frq^Δ733-774^, frq^Δ775-800^*, and *frq^Δ801-856^*. The data support that multiple regions of FRQ take part in the rhythmicity control and period determination as documented by a large body of literature.

**Figure 1.**
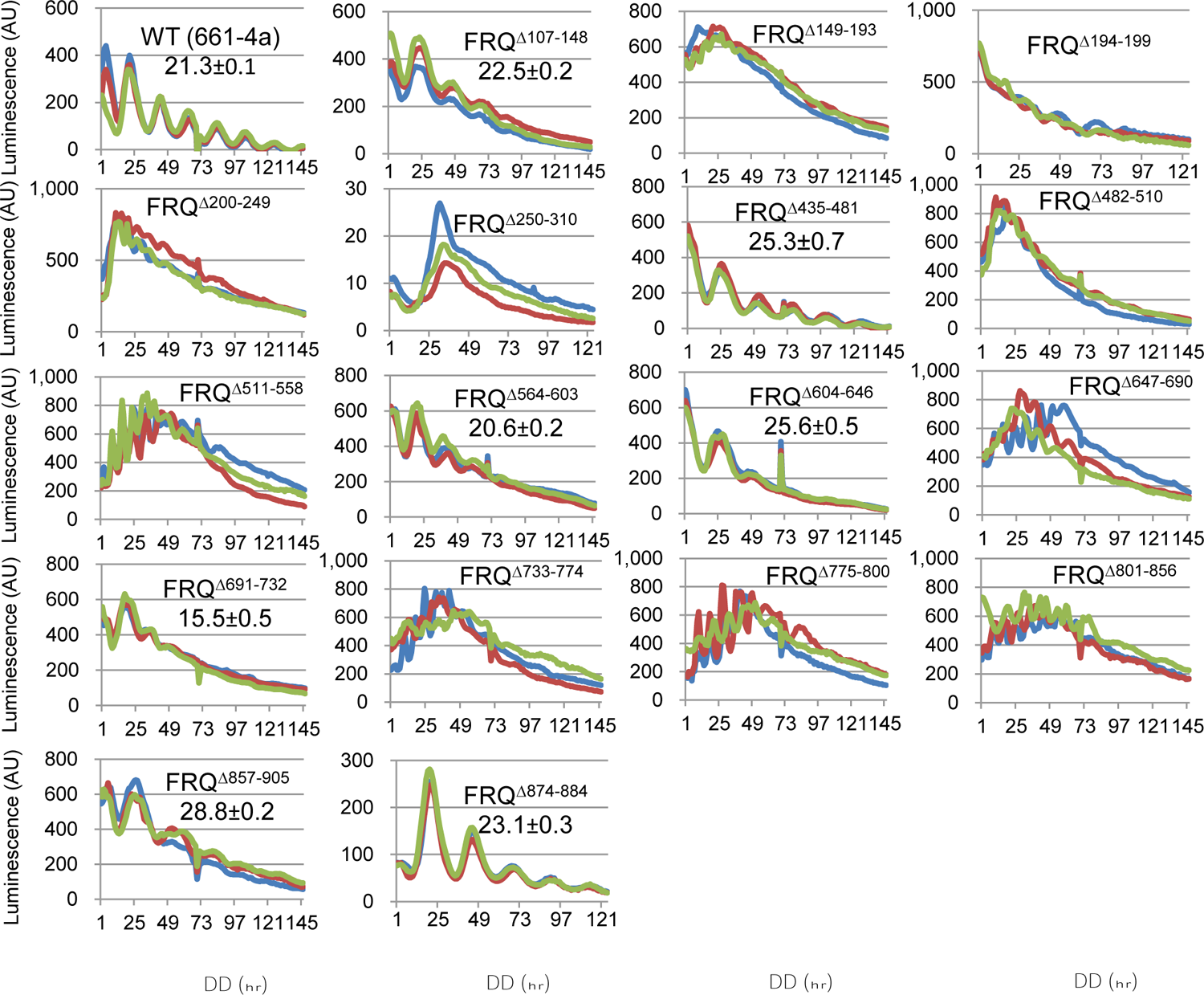
Luciferase assays of *frq* partial-deletion mutants at 25 °C in the dark. Strains were synchronized at 25 °C in the light overnight, and bioluminescence signals were tracked every hour by a CCD (charge-coupled device) camera after moving the strains to the dark at the same temperature. Three replicates (lines in different colors) were plotted for each mutant with the x-axis and y-axis representing time (in hours) and arbitrary units of the signal intensity, respectively. In this and subsequent figures, period length was calculated from three or more replicates and reported as the average +/- the standard error of the mean (SEM). All frq mutants throughout this study were targeted to its native locus with a tandem V5 and 6 x histidine (V5H6) tag at their C-termini.

### FRQ-WCC integrity is broken in arrhythmic strains

To examine the FFC-WCC formation that functions as the initial step in the repression elicited by FFC on WCC, FRQ (tagged with V5H6) in the frq mutants (in Figure 1) was pulled down with V5 resin, and FRQ along with FRH, WC-1, and WC-2 were blotted with protein-specific antibodies (Figure 2). When FRQ was enriched by immunoprecipitation in *frq^Δ149-193^, frq^Δ194-199^, frq^Δ200-249^, frq^Δ482-510^, frq^Δ511-558^, frq^Δ647-690^, frq^Δ733-774^, frq^Δ775-800^, frq^Δ801-856^*, and *frq^Δ857-905^*, WC-1 and WC-2 became undetectable with respect to the wild-type, and all except frq^Δ857-905^ showed an arrhythmic clock and enhanced C-box-driven reporter activity (Figures 1, 2). *frq^Δ107-148^, frq^Δ564-603^, frq^Δ604-646^, frq^Δ691-732^*, and *frq^Δ857-905^* displayed weak FFC and WCC interaction but sustained weak rhythmicity (Figures 1, 2). This result confirms that multiple domains on FRQ contribute cooperatively to contact with WCC. FRH dissociates from FRQ in *frq^Δ733-774^, frq^Δ775-800^*, and *frq^Δ801-856^* (Figure 2), which may lead to the loss of WCC in the complex as noted previously for *frq* mutants *frq^6B2^* and *frq^6B5^* (25). This also reinforces that the entirety of FRQ-FRH is needed for FFC to incorporate into WCC (7, 8, 25), presumably through constructing a proper quaternary structure of FFC. The clock in *frq^Δ733-774^, frq^Δ775-800^*, and *frq^Δ801-856^* does not oscillate (Figure 1), in line nicely with the finding that the absence of FRH in the FFC abrogates feedback loop closure (7, 8). *frq^Δ250-310^, frq^Δ435-481^*, and *frq^Δ874-884^* have normal FFC-WCC interaction (Figure 2). Collectively, the data here further characterize FRQ’s regions involved in WCC interaction but suggest that the binding strength of FFC-WCC is not indicative of the period length.

**Figure 2.**
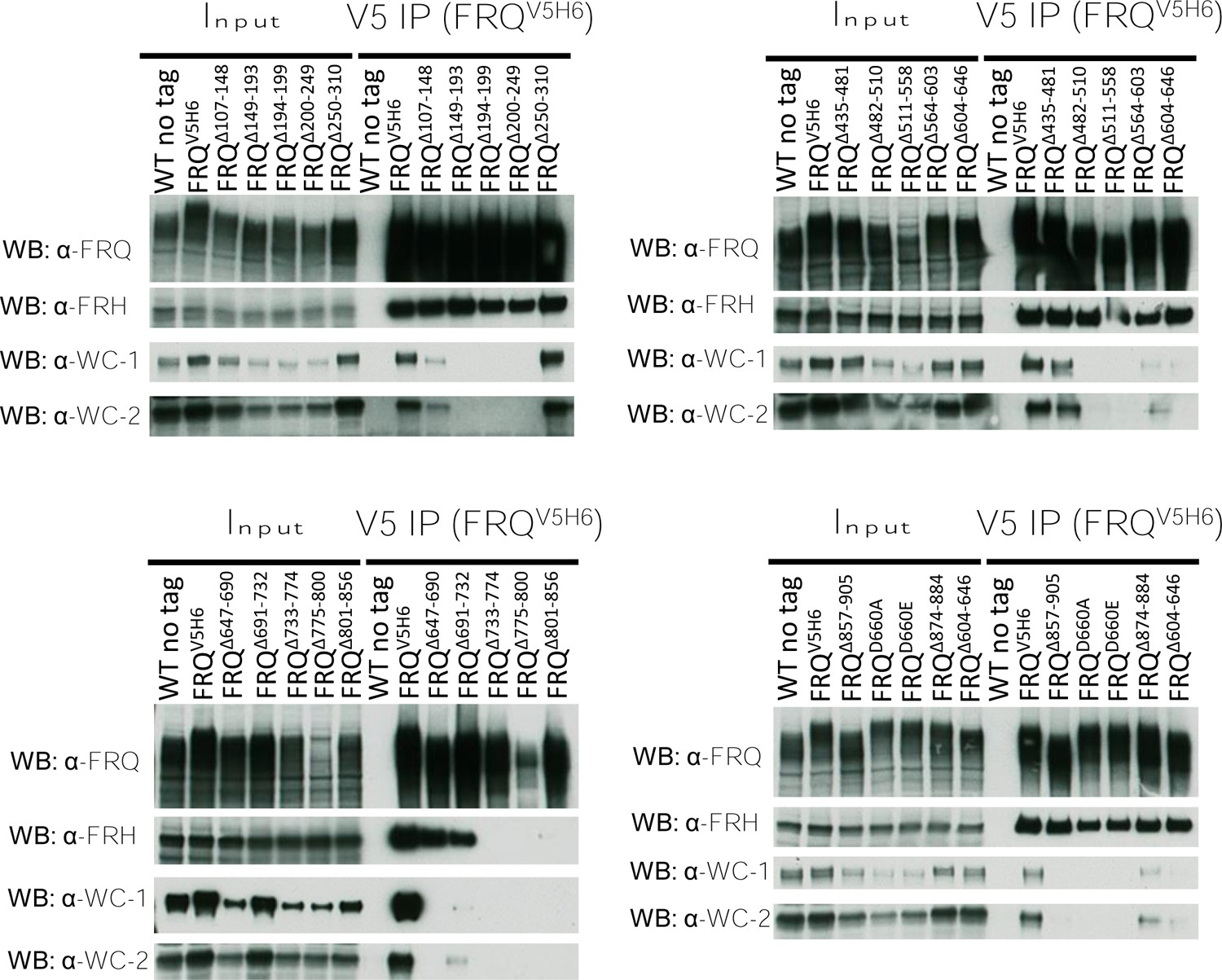
FRQ/FRH and WC-1/WC-2 interaction in the frq-partial-deletion mutants. FRQ (tagged with V5H6) was pulled down with V5 resin from a culture grown in constant light at 25 °C for 24 hrs, and western blotting (WB) was carried out using custom antibodies (raised from rabbits) against FRQ, FRH, WC-1, or WC-2 as indicated (see Materials and Methods for details). For V5 IP, WT (no tag) serves as the negative control, while WT FRQ tagged by V5H6 (FRQ^V5H6^) is the positive control. For “input”, 15-µg total protein was loaded per lane, and 10 µl out of a total of 100 µl eluate was loaded per lane for “V5 IP” samples. In this and subsequent WB-based analyses, comparable results were obtained in multiple biological replicate experiments.

### FFC-WCC interaction can be disrupted by high salt

A basic stretch (KKKRKRRK) near the Zinc-Finger-DNA-binding domain of WC-1 was previously identified as required for the WCC-FFC organization (20), which suggests that ionic charges may drive the complex formation. To confirm this hypothesis, the NaCl concentration in the regular protein-lysis buffer was raised from 137 mM that is used commonly in the biochemical analysis of FFC and WCC to 500 and 1,000 mM, either WC-1 or FRQ was immunoprecipitated, and the four clock components followed subsequently by Western blotting. Following WC-1 immunoprecipitation, the enrichment of WC-2 was barely affected by high salts in the buffer, but FRQ and FRH became wholly undetected at 500- or 1,000-mM sodium chloride (Figure 3A). Similarly, in a separate experiment with epitope-tagged FRQ instead, FRQ^V5H6^ as well as bound FRH were immunoprecipitated readily with V5 antibody regardless of salt titers, whereas WC-1 and WC-2 disappeared from the FFC when salt dosages rose (Figure 3B). The data support that FFC and WCC associate with each other by virtue of electrostatic forces, which are subject to high salt disruptions *in vitro*, while WC-1 and WC-2 bind together presumably by hydrophobic effects via their PAS domains (25); likewise, both FRQ and FRH possess specified domains buttressing their interaction (25, 32). Accordingly, interactions between FRQ and FRH as well as WC-1 and WC-2 are less prone to high ionic strength disruptions.

**Figure 3.**
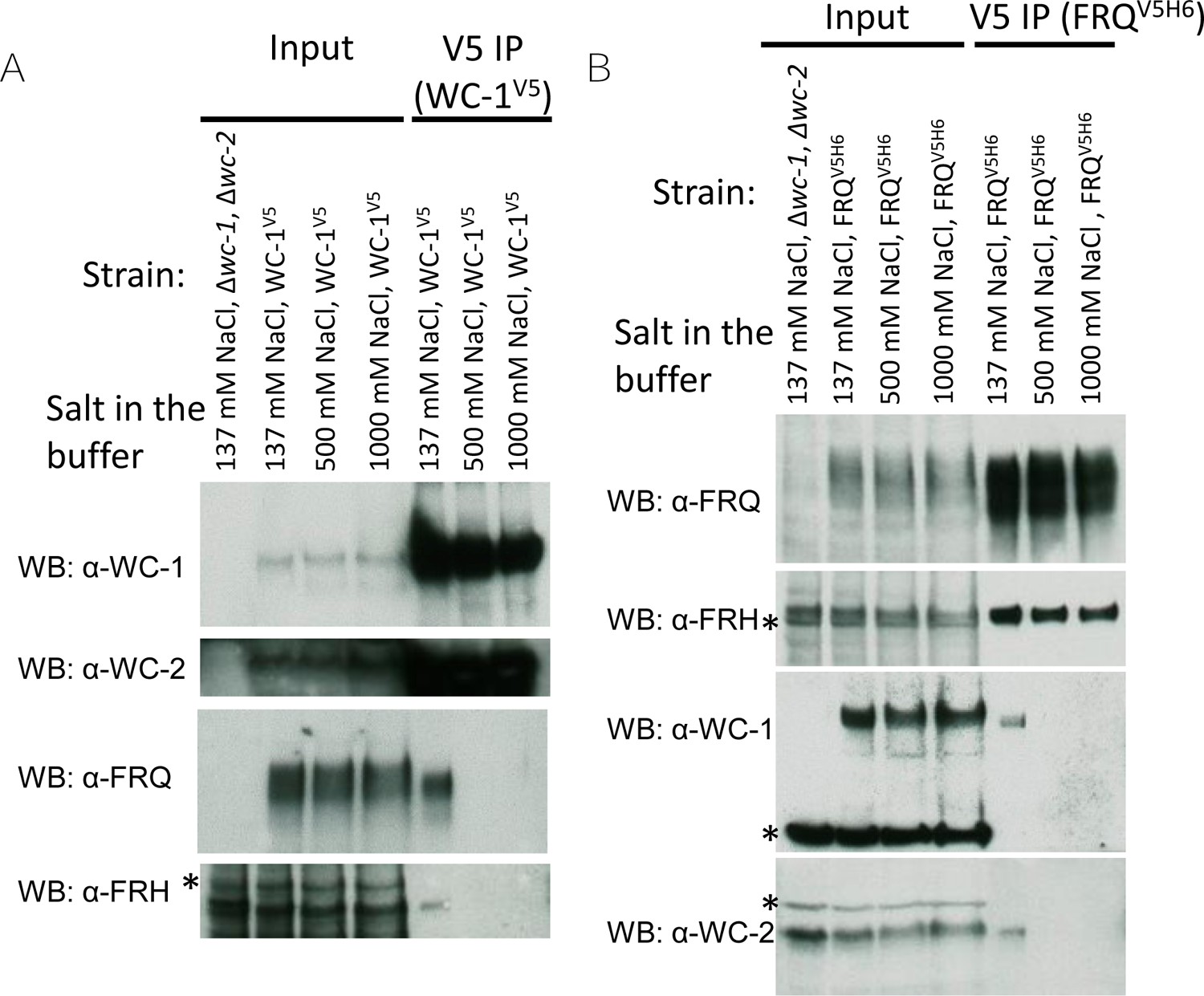
Increased salt concentration in the lysis/IP buffer disrupts the FFC-WCC complex. For IP (details can be found in Materials and Methods), the NaCl concentration in the lysis and wash buffers was set at 137, 500, or 1,000 mM as indicated above the panel. (A) WC-1^V5^ was cultured in the light at 25 °C for 24 hrs, harvested, and ground to a fine powder; buffers containing 137, 500, or 1,000 mM NaCl were added respectively to the ground powder for cell lysis; centrifugation-cleared lysate was incubated with V5 resin in a volume of one mL of the same lysis and wash buffer to pull down WC-1^V5^. Following IP, WB was carried out with WC-1-, WC-2-, FRQ-, or FRH-specific antibody as indicated. (B) The strain of FRQ^V5H6^ was grown at 25 °C plus light for ∼24 hrs, and FRQ^V5H6^ was pulled down with V5 resin. For both (A) and (B), 15-µg total protein was loaded per “input” lane, while each “V5 IP” lane contained 10 µl eluate out of a total of 100 µl. *Δwc-1, Δwc-2* serves as the negative control.

### Luciferase analysis of *frq* D/E-to-A mutants

Based on the observations that the organization of WCC-FFC is susceptible to the ambient ionic strength (Figure 3) and also that the basic motif of WC-1 is needed for the WCC-FFC connection (20), negatively charged residues, Asp (D)/Glu (E), in the key regions of FRQ (Figures 1 and 2) were mutated individually or in clusters to alanines for the purpose of pinning down FRQ’s residues involved in WCC binding. In subsequent luciferase reporter analyses, four classes of rhythm *alterations were noted in these frq D/E-to-A mutants (Figure 4): The clock became* fully arrhythmic in *frq^D149A, D150A, D156A, D157A^*, *frq^E200A, E202A, D207A^*, *frq^D738A, E739A, D740A, E747A, D748A^*, *frq^E760A, E763A, D773A, D774A^*, *frq^E801A, D802A, E805A, E809A^*, and *frq^E835A^*; period length was unaffected in *frq^D664A, D667A^*, *frq^E679A, E681A^*, *frq^D687A, E688A, E690A^*, *frq^D862A, D866A, D867A, D869A, D870A^*, *frq^D874A, D875A, E876A, E877A, E879A, E880A, E882A, E883A, D884A^*, *frq^E161A, E167A, E168A^*, and *frq^D243A^*; *frq^D738A, E739A, D740A, E747A, D748A^* and *frq^E888A, D901A^* display decreased period lengths of 16.8 and 19.6 hrs respectively; period length of the rhythms was elongated by ∼2, 3, 5, and 7 in frq^D214A, E217A^, frq^E552A, D553A, E555A, D556A^, frq^D532A, D539A, E542A, D543A^, and frq^E511A, D521A^ respectively. The data of period changes here agree fairly well with the period-alternating pattern of frq mutants from a recent publication demonstrating that removal of phosphosites in the N-terminal and middle regions of FRQ results typically in prolonged period lengths while elimination of C-terminal phosphorylations always lessens the period length (27).

**Figure 4.**
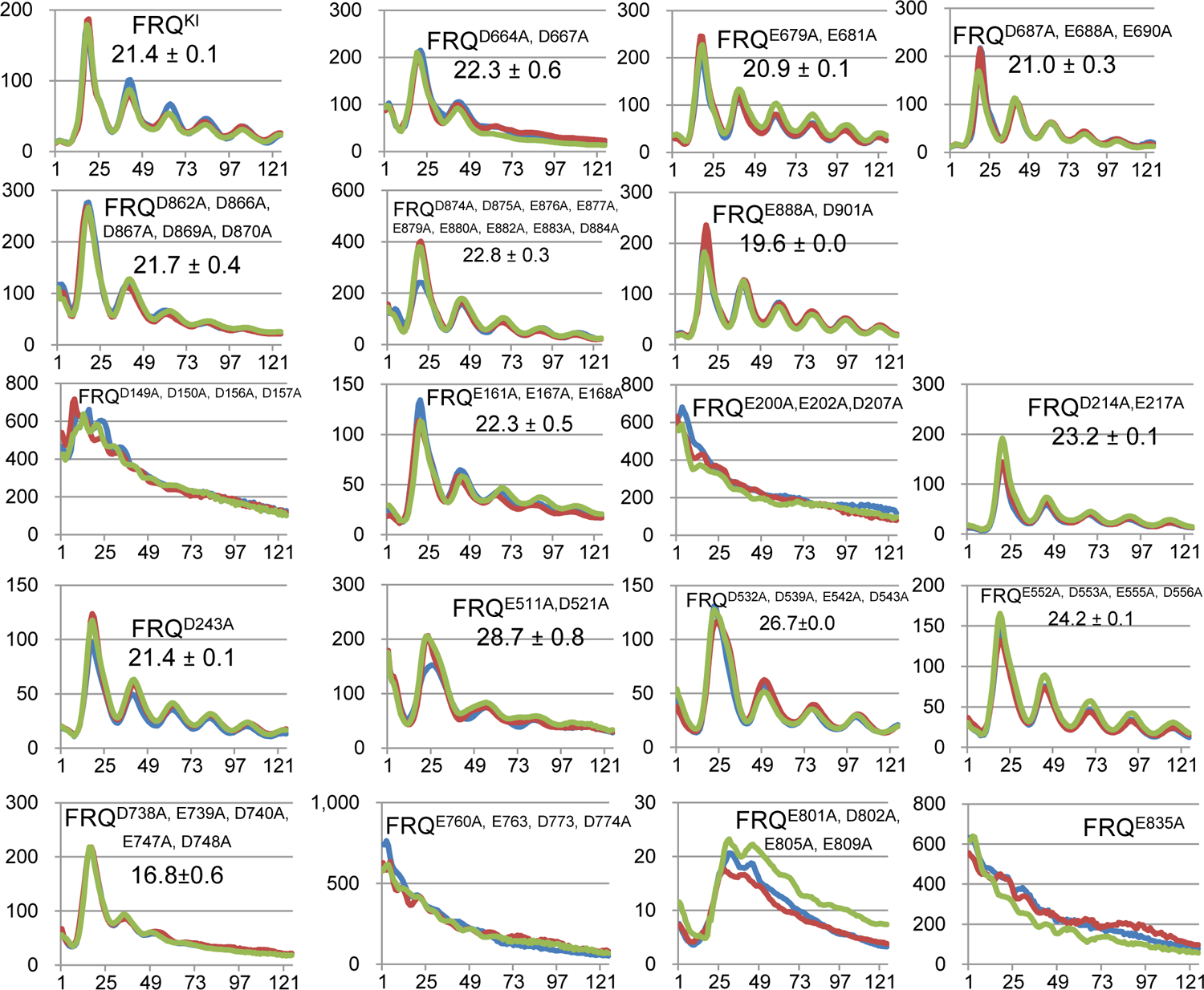
Luciferase assays of frq D/E-to-A mutants at 25 °C in the dark. Synchronization of strains was done in the light at 25 °C overnight, and light signals were tracked every hour by a CCD camera following the transfer of the strains to the dark at 25 °C. Lines in different colors represent three replicates with the x-axis and y-axis displaying time (in hours) and the signal intensity (arbitrary units), respectively. Period length was determined from three replicates and shown as the average +/- the SEM. All frq mutants bear a V5H6 tag at their C-termini, targeting at the frq native locus.

### The FFC-WCC establishment is impaired in certain frq D/E-to-A mutants

To determine whether the FFC-WCC in the D/E-to-A mutants from Figure 4 remains intact biochemically, we examined expression and interaction of the four core-clock components, FRQ, FRH, WC-1, and WC-2 by immunoprecipitation and Western blotting. FRQ was not detected in *frq^D149A, D150A, D156A, D157A^* and *frq^E835A^*, and was only very weakly detected in *frq^E760A, E763A, D773A, D774A^* after enrichment (Figure 5), suggesting that these four residues may significantly impact FRQ stability; this explains why they displayed arrhythmicity but high signal intensity in the luciferase analysis (Figure 4), basically mirroring the behavior of Δ*frq* or *frh* mutants (7, 8, 33). FRQ abundance in *frq^E200A, E202A, D207A^* drops drastically, which might lead to a reduction of FRH in the complex. The little to no expression of FRQ along with diminished abundance of WCC in *frq^D149A, D150A, D156A, D157A^*, *frq^E200A, E202A, D207A^*, *frq^E760A, E763A, D773A, D774A^*, and *frq^E835A^* is nicely compatible with the “black widow model” proposing a negative correlation between the activity of transcription factors in transcription with their cellular abundance (8, 10, 34).

**Figure 5.**
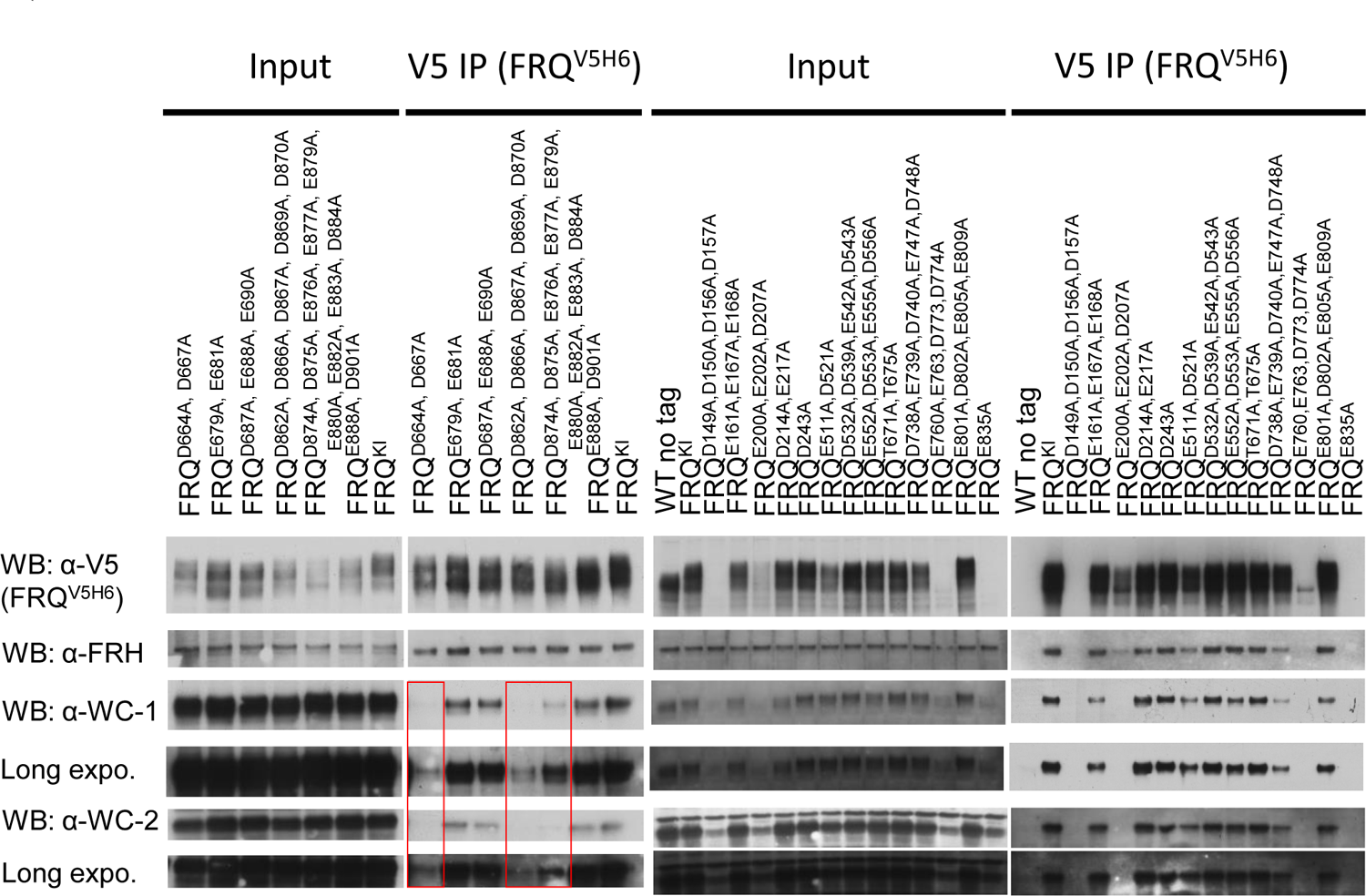
Two clusters of D/E in the C-terminus of FRQ are essential for associating with WCC. FRQ (bearing a V5H6 tag) was pulled down with V5 resin from a culture grown in constant light at 25 °C, and WB was performed using antibodies against FRQ, FRH, WC-1, or WC-2 as indicated. WT FRQ tagged by V5H6 (FRQV5H6) is the positive control in the assay. Red boxes mark the absence of or an extremely reduced amount of FFC-bound WCC.

Interestingly, WC-1 and WC-2 levels in the FFC-WCC in frq^D664A, D667A^, frq^D862A, D866A, D867A, D869A, D870A^, *frq^D874A, D875A, E876A, E877A, E879A, E880A, E882A, E883A, D884A^* become dramatically reduced (can only be visualized with a long exposure in Western blotting) in relation to WT, though FRQ and FRH in these strains interact normally with each other (Figure 5). This result appears to be astonishing as the three frq mutants demonstrate an approximate WT period (Figure 4), but it evidently suggests that the interaction strength between the positive and negative element complexes does not predict the period length, and also suggests that FRQ-promoted phosphorylation of WCC can persist efficiently even with less WCC bound in the complex.

### The third cluster of D/E on FRQ crucial for the WCC-FFC formation

FRQ was completely undetectable in *frq^D149A, D150A, D156A, D157A^, frq^E760A, E763, D773A, D774A^*, and *frq^E835A^* (Figure 5). To probe the role of these D/E clusters, we made frq mutants bearing fewer mutations to these sites, or an E835D substitution, and monitored C-box activity and WCC-FFC establishment. FRQ’s E183 and E187 were also included in the mutagenesis because they are located nearby D149, D150, D156, and D157. The clock in *frq^D149A, D150A^, frq^E760A, E763A^*, and *frq^E835D^* oscillates robustly with an approximately WT period, while *frq^D156A, D157A^, frq^E183A, E187A^*, and *frq^D773A, D774A^* completely lost rhythmicity (*frq^E183A, E187A^* only shows one peak) (Figure 6A). FRQ expression in *frq^D773A, D774A^* is extraordinarily low (a faint band after immunoprecipitation) (Figure 6B), explaining the arrhythmicity in the luciferase assay (Figure 6A). WC-1 and WC-2 levels in FFC-WCC became greatly diminished in *frq^D149A, D150A^*, once again showing that the WCC-FFC abundance in the cell does not faithfully reflect the period length (Figure 5). The loss of FFC-WCC in *frq^E183A, E187A^* and *frq^D773A, D774A^* (Figure 6B) matches their utterly abolished rhythmicity (Figure 6A). The acquired data here reinforces the necessity of incorporating FFC into WCC for eliciting phosphorylation-induced repression. It is noteworthy that unlike frq^E835A^ in which FRQ protein accumulation was abolished (Figure 5), *frq^E835D^* bears a vigorously oscillating clock as well as normal FRQ abundance and unchanged interaction of FRQ with WCC and FRH in comparison with WT (Figure 6B), suggesting that the negative charge of E835 is the main contributor in promoting FRQ accumulation.

**Figure 6.**
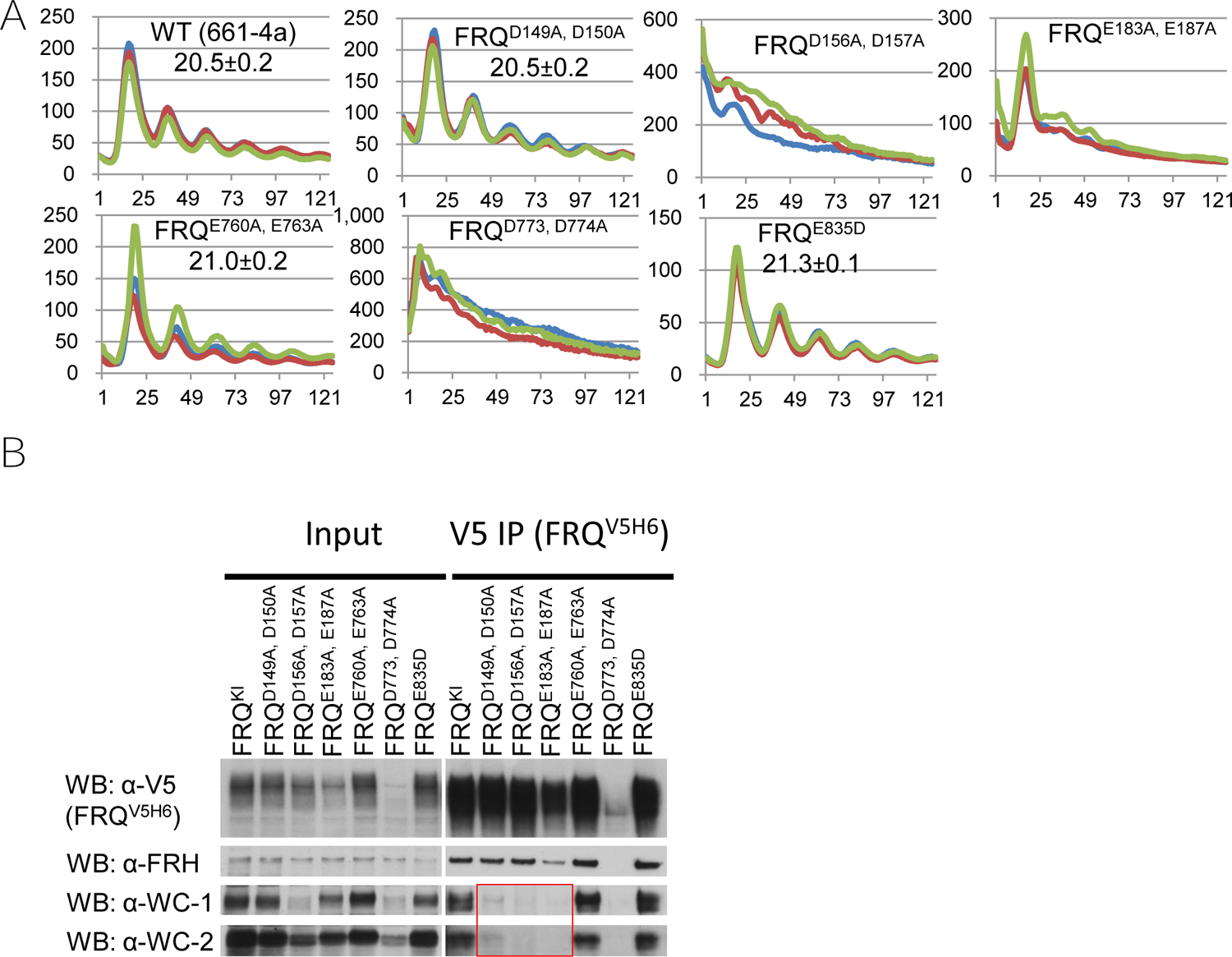
FRQ residues D149, D150, D156, D157, E183, E187 are needed for FFC to bind WCC. (A) Luciferase assays of indicated frq mutants at 25 °C in the dark. Synchronization for the clock was carried out at 25 °C plus light overnight, and then bioluminescence signals from the darkness-grown strains (at 25 °C) were tracked every hour. Three replicates are represented by differently colored lines with time (in hours) as the x-axis and the signal intensity (arbitrary units) as the y-axis. Period length was measured from three replicates and displayed as the average +/- the SEM. All frq alleles were appended by a V5H6 tag at their C-termini at the frq locus. (B) Interaction of clock components FRQ, FRH, WC-1, and WC-2 in the stated frq mutants. V5H6-tagged FRQ was immunoprecipitated with V5 resin from strains cultured in the light at 25 °C for ∼24 hrs, and WB was carried out with indicated antibodies against FRQ, FRH, WC-1, or WC-2 (see Materials and Methods for details). The red box denotes remarkably decreased or totally undetected WCC from the FRQ pull-down (by V5).

### Expression and clock-components interaction in *frq* truncations

Certain residues on FRQ, like D773, D774, and E835, contribute significantly to the accumulation of the full-length protein (Figures 5 and 6). We wondered whether this observation also applies to truncated variants of FRQ and whether retaining certain clusters of important D/E suffices for WCC recruitment. Hence, we made two set of frq mutants each bearing abridged versions of FRQ and together covering the whole FRQ twice: *frq^1-558^ and frq^559-989^, frq^1-310^, frq^311-628^*, and *frq^629-989^* (Figure 7A). As expected, *frq^1-310^, frq^311-628^*, and *frq^629-989^* totally lost rhythmicity in the luciferase assay (Figure 7A). The FRQ level in *frq^1-310^* fell below our detection limit in WB even after an enrichment by immunoprecipitation, while in other mutants it appears to be comparable to that in WT (Figure 7B). Although FRH complexed with FRQ as strongly in *frq^559-989^* and *frq^629-989^* as in WT, WC-1 and WC-2 in all of these mutants was undetectable above the background in WB (Figure 7B), indicating that preservation of individual D/E clusters is not sufficient for FFC-WCC formation.

**Figure 7.**
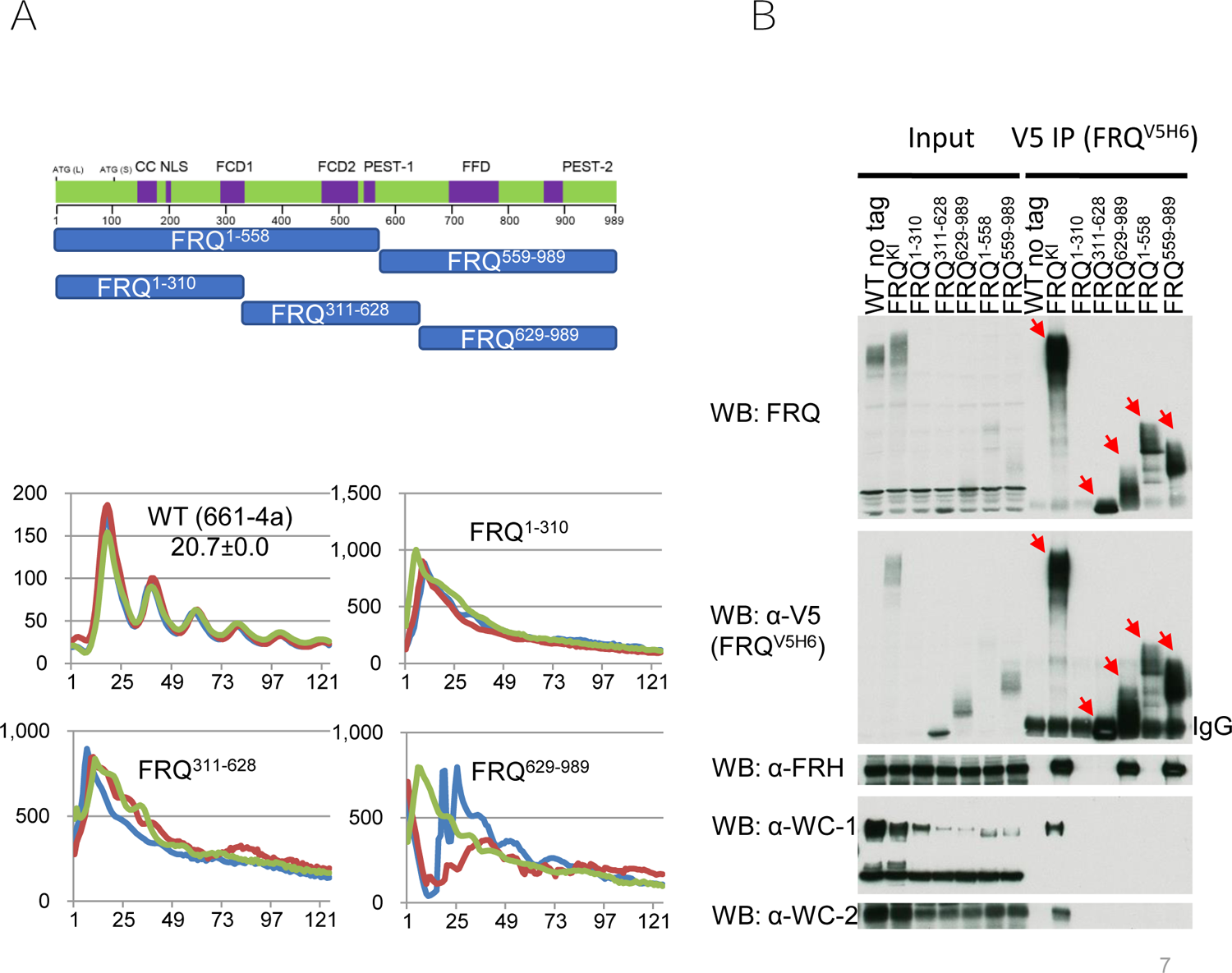
Truncated variants of FRQ have variable expression levels and WCC&FRH interactions. (A) Schematics of frq mutants covering the whole FRQ open-reading-frame. Upper, each blue bar means a frq mutant encoding a polypeptide with the beginning and ending residues labelled within the bar. All frq mutants here carry a V5H6 at the C-terminus just as others in this study. Bottom, luciferase analysis of FRQ^1-310^, FRQ^311-628^, and FRQ^629-989^ each carrying a C-box promoter-driven luciferase gene at 25 °C in the dark. (B) Expression and interaction of FRQ, FRH, WC-1, and WC-2 in the indicated frq mutants. Immunoprecipitation was done with V5 resin to pull down V5H6-tagged FRQ from strains cultured in the light at 25 °C for ∼24 hrs, and WB was carried out with indicated antibodies against FRQ, FRH, WC-1, or WC-2. Red arrows point to FRQ bands in V5 immunoprecipitations. Note: The FRQ band in *frq^1-310^* partially overlaps with the IgG band in the WB of V5 but is evident in the top blot with FRQ-specific antibody.

### Glu (E) substitution of FRQ’s D660 phenocopies *frq^D660A^*

FRQ’s residue D660 was found to be essential for FFC to recruit WCC and thereby for the negative feedback loop (35). To test whether retaining the negative charge of D660 justifies the role of FRQ in the clock, we generated *frq^D660E^* and assayed it for C-box activity by luciferase analysis and WCC interaction by immunoprecipitation. To our surprise, *frq^D660E^* behaved as arrhythmically as *frq^D660A^* (Figure 7A) and failed to rescue the loss of the FFC-WCC in *frq^D660A^* (Figure 1). The data reveal that the negative charge of D660 is not the sole factor in determining the FFC-WCC formation and circadian rhythmicity, an observation that contrasts sharply with what we have obtained from *frq^E835A^* and *frq^E835D^* (Figures 5 and 6), demonstrating the importance of the side-chain charge of E835 in promoting FRQ accumulation.

## Discussion

In *Neurospora*, the *frequency* gene encodes the central scaffolding protein FRQ that organizes the multicomponent negative arm of the core clock, and the cellular timing information has been encoded sophisticatedly and situated precisely in its expression and probably more importantly in its chemical modifications. Unfortunately, the complete FRQ structure, like that of its mammalian counterparts the PERs, remains unresolved due mainly to its scarcity and intrinsically disordered nature. This leaves many fascinating questions concerning the core-clock operation unsettled. For instance, how do the negative arm complexes FFC and PER/CRY accurately bridge kinases to progressively inhibit WCC or CLOCK/BMAL1 via phosphorylation in the repressive phase of the clock; how is it that FFC only binds to a small percentage of WCC but is able to proficiently repress the whole WCC pool. In this study, we attempted to probe these queries by searching for important regions and more specifically key residues of FRQ engaging in WCC contacts. Segment-deletion analysis identified regions on FRQ required for the core oscillator as well as WCC recruitment (Figures 1 and 2). In agreement with prior literature, more than one region on FRQ contributes to WCC interaction (Figure 2). Biochemical analysis further revealed that the FFC-WCC complex is vulnerable to elevated salt; the DBD on WC-1 was identified as a key for FFC interaction. Altogether, these leads guided us to dissect negatively charged residues falling in the regions of FRQ that were found to be important for WCC interaction (Figure 1). In addition, we unintentionally noted residues vital for FRQ accumulation or stability as well as the ones significantly impacting period length when mutated (Figure 4). Immunoprecipitation assays confirmed that three clusters of negatively charged amino acids, D149/D150/D156/D157/E183/E187, D664/D667, and D/E from aa 862-D884 (Figure 8B), are required for WCC association, but surprisingly, *frq^D664A, D667A^*, *frq^D862A, D866A, D867A, D869A, D870A^*, *frq^D874A, D875A, E876A, E877A, E879A, E880A, E882A, E883A, D884A^* displayed vigorous circadian rhythms with a period length almost identical to WT. These data indicate that the interacting quality between the positive and negative elements does not control the pace of the oscillator. It remains elusive whether these FRQ’s D/E residues discovered in this study are located on the interface of the FRQ structure for contacting directly with WCC or they contribute merely to the maintenance of FRQ’s tertiary structures that are needed in WCC recognition.

**Figure 8.**
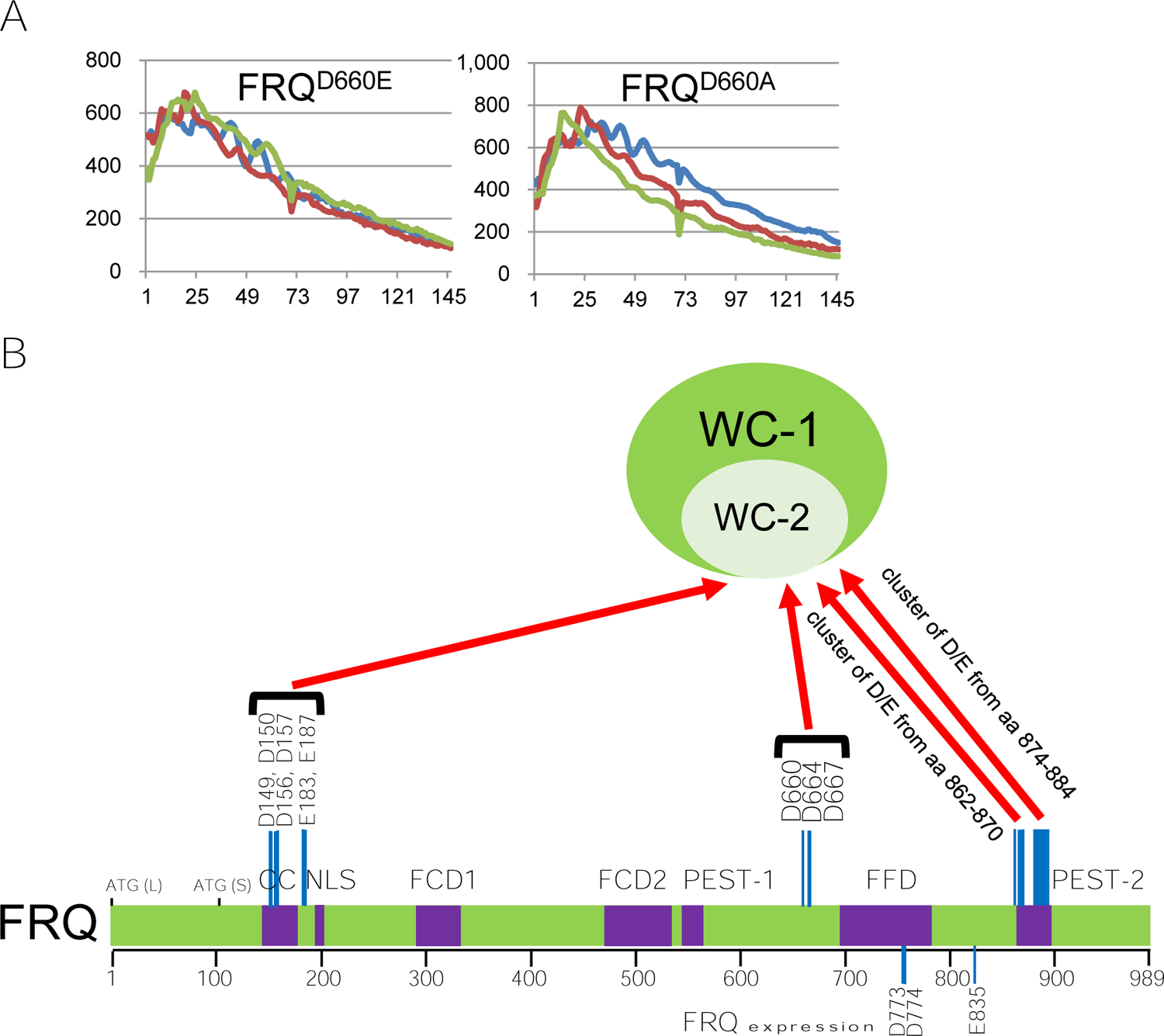
A working model illustrating FRQ’s D/E clusters that mediate the WCC-FFC formation. (A) Not all FCC-WCC interactions are electrostatic. Luciferase analysis of FRQ^D660A^ and FRQ^D660E^. Data here are presented in a similar manner as in prior figures with x and y axes representing hrs and arbitrary units of the luminescence intensity (B) A depiction summarizing the negatively charged residues (D and E) on FRQ in controlling WC-1 and WC-2 interaction. The cluster of D/E from amino acids 862-870 contains D862, D866, D867, D869, and D870; that from aa 874-884 includes residues D874, D875, E876, E877, E879, E880, E882, E883, and D884. The red arrows represent that these residues in FRQ contribute either directly (through physical contacts) or indirectly (such as constructing or maintaining regional structures) to binding WCC.

Recently, the binding strength of FRQ-CKI has been demonstrated to be an indicator of the period length at normal temperatures as well as in a physiological temperature range (36, 37). The correlation of WCC-FRQ abundance with period length is vague since *frq* mutants with a constant WCC-FRQ level showed varying period lengths (36). Most WCC in the cell resides in the nucleus, while the majority of FRQ is located in the cytoplasm (11, 20, 23, 31). These observations underpin the fact that even in the WT strain, only a small portion of the FFC pool encounters and then complexes with WCC (11). The FFC-WCC super-complex (in terms of size) (11) may assemble in a dynamic manner as evidenced by the quantitative proteomic data showing that WCC interacts preferentially with hypophosphorylated FRQ, whereas FRH associates constantly with FRQ independent of the latter’s phosphorylation status. Unexpectedly, we here saw that the period length in certain *frq* mutants was almost unaffected even with the severely impaired WCC-FFC establishment. Based upon prior evidence, this may be interpreted in several ways. First of all, there may be only a limited fraction of the WCC pool actively participating in *frq* transcription, and correspondingly timely expressed FRQ only needs to repress this small amount of DNA-bound WCC. Thus, the core oscillator could operate in a normal way even in mutants composed of remarkably less FRQ and WCC (similar to the scenario in WT with most FFC and WCC not participating in the core oscillator). This has been suggested by experimental data showing that a manifest downregulation of WCC does not greatly perturb circadian rhythmicity beyond a slight ∼ 2 hrs period lengthening (34). Second, the dynamic organization of WCC-FFC may ensure that the clock can be sustained with less WCC and FFC involved. Previous findings suggested that both WCC and FRQ rapidly translocate between the nucleus and cytoplasm (38, 39). Finally, the course of FFC-mediated WCC repression via phosphorylation lasts more than roughly half a day (40, 41). Therefore, FFC in certain mutants may be still fully capable of bringing about thorough inactivation to WCC over a long while even though the efficiency may be not as high as that in WT. Apparently, these possibilities are not mutually exclusive. Future investigations on these uncertainties would benefit our deeper understanding of the fundamental properties of the molecular clock.

FRQ phosphorylation has been extensively explored for decades as the primary regulatory mechanism governing the *Neurospora* clock (27–30, 42, 43, 43–49). Freshly translated hypophosphorylated FRQ possesses higher affinity for WCC than the massively phosphorylated isoforms found at subsequent circadian times, implying that multisite phosphorylation events on FRQ modulate the FFC-WCC assembly. Phosphorylation of FRQ may negatively impact WCC’s accessibility to certain important D/E residues of FRQ. Alternatively, mutations to these D/E residues may impinge on nearby phosphoevents, although we did not notice obvious alterations of the overall phosphorylation profiles of FRQ in these mutants (Figures 5 and 6B). Needless to say, future research is desirable to solve these intriguing mysteries.

## Materials and methods

### Strains and growth conditions

*Neurospora* strain 661-4a (*ras-1^bd^, A, his-3::C-box-driven luciferase*), which serves as WT in luciferase assays, contains the *frq C-box* promoter fused to the codon-optimized firefly *luciferase* (transcriptional fusion) at the *his*-3 locus (40, 50).

### Growth conditions

All vegetative cultures were maintained on complete-medium slants bearing 1 x Vogel’s, 1.6% glycerol, 0.025% casein hydrolysate, 0.5% yeast extract, 0.5% malt extract, and 1.5% agar (51). *Neurospora* sexual crosses were performed on Westergaard’s agar plates containing 1 x Westergaard’s salts, 2% sucrose, 50 ng/mL biotin, and 1.5% agar (52).

### *frq* mutant generation

*frq* mutants were created by a modified method using yeast homologous recombination-based integration of PCR fragments (27): Restriction-digested shuttle vector *pCB05* (30) was recombined with PCR (Thermo Fisher Scientific, Catalog # F549S) products amplified with primers bearing partial deletions or point mutations of FRQ. Four primer pairs worked as flanks in homologous recombination in a yeast strain (FY834). To introduce deletions falling in aa 1-214 of FRQ, two PCR reactions were performed: one with a forward primer “*frq* segment 1F” (5’-GAACCAGAACGTAGCAGTGTG-3’) and a reverse primer “del.aa # to # R” bearing a deletion to FRQ and the other using a forward primer “del.aa # to # F” which is reverse and complementary to “del.aa # to # R” and a reverse primer “*frq* segment 1R” (5’-GACGATGACGACGAATCGTG-3’), and then the two PCR products were co-transformed into yeast along with *pCB05* digested with *Bst*XI (New England Biolabs [NEB], Catalog # R0113S) and XhoI (NEB, Catalog # R0146S) to generate a circular construct. Likewise, for mutations falling in aa 215-437 of FRQ, primers “*frq* segment 2F” (5’-GTGAGTTGGAGGCAACGCTC-3’) and “*frq* segment 2R” (5’-GTCCATATTCTCGGATGGTA-3’ were used for PCRs in combination with *pCB05* digested with *XhoI* (NEB, Catalog # R0146S) to *NruI* (NEB, Catalog # R0192S); “*frq* segment 3F” (5’-GTCGCACTGGTAACAACACCTC-3’) and “*frq* segment 3R” (5’-CAGCACATGTTCAACTTCATCAC-3’) were designed for *pCB05* digested with NruI (NEB, Catalog # R0192S) and *FseI* (NEB, Catalog # R0588S) (FRQ aa 438-675), and “*frq* segment 4F” (5’-CACCGATCTTTCAGGAGACCCTG-3’) and “*frq* segment 4R” (5’-CACTCAGGTC TCAATGGTGA TG-3’) pair with *pCB05* digested with FseI (NEB, Catalog # R0588S) and *MluI* (NEB, Catalog # R0198S) (FRQ aa 676-989). If deleting regions or mutating residues involve two or more segments above, corresponding restriction enzymes and primers encompassing the regions were selected and combined for yeast recombination. All targeted mutations were validated by cycle sequencing with *frq*-specific primers at the Dartmouth Core facility. All these *frq* variants were targeted for homologous recombination at its native locus. Plasmids verified by Sanger sequencing were linearized with AseI (NEB, Catalog # R0526S) and *SspI* (NEB, Catalog # R0132S) and purified with the QIAquick PCR Purification Kit (Qiagen, Catalog # 28104) for *Neurospora* transformation. *Neurospora* transformation via electroporation (settings: 1,500 V, 600 Ω, and 25 μF) was performed using an electroporator (BTX, Model # ECM 630) as previously reported (53). The recipient strain for generating *frq* mutants is *Δfrq::hph; Δmus-52::hph; ras-1^bd^; C-box luciferase* at *his*-3, and all *frq* mutants made in this research bear the ras-1^bd^ mutation (54) and *frq C-box-driven* codon-optimized firefly *luciferase* gene at the *his*-3 locus for phenotype analyses, and they also contain a V5H6 tag at their C termini for biochemical assays.

### Protein lysate and Western blot

Protein lysates for Western blots (WB) were prepared as previously described (55). Liquid medium (LCM) for culturing *Neurospora* is composed of 1 x Vogel’s, 0.5% arginine, 50 ng/mL biotin, and 2% glucose. Vacuum-dehydrated *Neurospora* tissue was frozen thoroughly in liquid nitrogen and ground to a fine powder using a mortar and pestle, the protein-extraction buffer (50 mM HEPES [pH 7.4], 137 mM NaCl, 10% glycerol, 0.4% NP-40) with cOmplete, Mini, EDTA-free Protease Inhibitor Cocktail (Roche, Catalog # 04693159001, at a dilution of 1 tablet to 10 mL buffer) was added to the powder, and the mixture of the *Neurospora* tissue powder and buffer was treated with cycles of vortexing for 10 sec and resting on ice for another 10 sec for a total of 2 min. For WB, equal amounts (15 μg) of centrifugation (12,000 rpm at 4 °C for 10 min)-cleared whole-cell lysate were loaded per lane in a commercial 3-8% 1.5-mm x 15-well Tris-Acetate SDS gel (Thermo Fisher Scientific, Catalog # EA03785BOX) with 1 x NuPAGE Tris-Acetate SDS Running Buffer (Thermo Fisher Scientific, Catalog # LA0041). Rabbit V5 antibody (Abcam, Catalog # ab9116), mouse FLAG antibody (Sigma-Aldrich, Catalog # F3165), or rabbit HA (Abcam, Catalog # ab9110) were diluted at 1:5,000 as the primary antibodies (47). Custom rabbit FRQ, FRH, WC-1, and WC-2 antibodies have been described previously for applications in WB (6, 56, 57).

### Immunoprecipitation (IP)

IP with *Neurospora* lysate was performed as previously described (20). Briefly, 2 mgs of total protein (cleaned by 12,000-rpm centrifugation at 4 °C for 10 min) were incubated with 20 μL of V5 agarose (Sigma-Aldrich, Catalog #7345) by rotating at 4 °C for 2 hrs. The agarose beads were then washed twice with the same protein extraction buffer (50 mM HEPES [pH 7.4], 137 mM NaCl, 10% glycerol, 0.4% NP-40) and eluted by adding 100 µL of 5 × SDS sample buffer and then being heated at 99 °C for 5 min. 10 out of the 100 µL IP were loaded per lane in WB.

### *Luciferase* assays

*Luciferase* assays were performed as previously described (58, 59). 96-well plates with each well containing 0.8 mL of the *luciferase*-assay medium were inoculated with conidial suspension, and the inoculated strains were grown at 25 °C plus constant light for 16–24 hrs and then transferred to the dark at the same temperature for recording the light production. Bioluminescence signals were recorded with a CCD camera every hour, data were obtained with ImageJ and a custom macro, and period lengths of the strains were manually calculated. Raw data from three replicates were shown in the figures, and time (in hours) is on the x-axis while arbitrary units of the signal intensity is on the y-axis. Medium for *luciferase* assays contains 1 x Vogel’s salts, 0.17% arginine, 1.5% bacto-agar, 50 ng/mL biotin, 0.1% glucose, and 12.5 μM luciferin (GoldBio, Catalog # LUCK-2G). WT used in *luciferase* assays was 661-4a (ras-1^bd^, A) containing the *frq* C-box fused to the codon-optimized firefly *luciferase* gene (transcriptional fusion) at the *his-3* locus.

## Acknowledgements

This work was funded by the National Institutes of Health to Jay C. Dunlap (R35GM118021).

## Data availability

The *Neurospora* strains made in this study are available upon request. All data used to draw conclusions of the article have been presented within the figures.

